# Childhood conditions, pathways to entertainment work and current practices of female entertainment workers in Cambodia: Baseline findings from the Mobile Link trial

**DOI:** 10.1101/618983

**Authors:** Carinne Brody, Pheak Chhoun, Sovannary Tuot, Dallas Swendeman, Siyan Yi

**Affiliations:** Public Health Program, College of Education and Health Sciences, Touro University California, Vallejo, CA, the United States; KHANA Center for Population Health Research, Phnom Penh, Cambodia; Department of Psychiatry and Biobehavioral Sciences, University of California, Los Angeles, CA, the United States; Saw Swee Hock School of Public Health, National University of Singapore and National University Health System, Singapore, Singapore

**Keywords:** Entertainment work, gender-based violence, HIV, human rights, sexual and reproductive health, developing country

## Abstract

**Background:** Entertainment venues have been identified as an important location for HIV prevention due to the increasing number of young female entertainment and sex workers at these venues. The purpose of this report is to increase understanding of the childhood conditions, pathways to entertainment work and current practices of female entertainment workers (FEWs) in Cambodia.

**Methods:** Data used for this study were collected in April 2018 as part of the baseline survey of the Mobile Link, a randomized controlled trial to improve sexual and reproductive health of FEWs in Cambodia. We used a stratified random sampling method to recruit 600 FEWs for face-to-face interviews using a structured questionnaire. Descriptive analyses were performed.

**Results:** Most participants came from childhood homes without electricity (82.0%) or running water (87.0%). Most women moved to the city in the last ten years (80.5%) for economic reasons (43.7%). About a third worked in the garment industry prior to the entertainment industry (36.7%). Participation in transactional sex in the past three months was reported by 36.0%. Women reported low condom use practices with non-paying partners (23.4% used condom with partner at last sex), excessive and forced alcohol use at work (33.1% reported being forced to drink alcohol at work more than once a month), low modern contraception use (31.4% was using modern contraception), and experiences of gender-based violence (23.3% reported verbal threats, physical abuse or forced sex in the past six months).

**Conclusions:** This information will help to support the development of future individual and structural level interventions for the safety and support of FEWs. In addition, these results may contribute to an evidence base that can inform policy level changes intended to support the realization of full human rights for entertainment works in Cambodia including the rights to health, safety, and respectful employment.

## Introduction

In Cambodia, as in many parts of Asia, one common pathway to a livelihood for young women from poor families is to migrate to urban areas to earn a better wage and send money to families [1]. Many young women in Cambodia work in garment factories, which are the backbone of Cambodia’s economy producing over 70% of Cambodia’s total exports and employing over 700,000 people [2], who are typically women who begin working in factories as teens despite child labor laws [3]. Over 65% of garment factory workers are under 24 years old [4]. They receive low pay, work long hours, and can struggle to navigate new social norms away from family oversight [5] and their former social support networks [6]. As a result of the poor wages, many young women seek to supplement or change to better paying jobs at entertainment venues such as beer gardens, massage parlors, and karaoke bars. In these roles, many women will engage in entertainment work, which can include transactional sex [2,7,8]. In 2014, the estimated number of female entertainment workers in Cambodia was approximately 40,000 [9], and by 2019, it was around 70,000 [10].

Entertainment work can include risks to women’s health, safety, and economic survival, and the industry has been a target of HIV prevention campaigns. Women account for over half (52%) of all HIV infections in Cambodia, higher than the regional average (35%) [11]. The government of Cambodia is calling for urgent preventative action targeting young “at-risk” women [2]. Entertainment venues have been identified as an important location for the prevention of HIV and other sexually transmitted infections (STIs) due to the increasing risk along a pathway from rural-urban migrants to factory workers to entertainment workers and sex workers [1]. While community outreach, HIV peer education, and place-based testing is now available for female entertainment workers in Cambodia, recent studies have found that only 53% reported a recent HIV test (in the past six months) [12], and 31% reported consistent condom use with romantic partners [13]. While 45% reported using modern contraceptives, 21.4% reported having at least one induced abortion since starting work in entertainment venues [14]. In addition, mental health indicators in this population are concerning, with 43% reporting a high level of psychological distress [15].

The mental and physical health of female entertainment workers intersects with their occupational health and safety in entertainment venues. Occupational safety concerns reported by female entertainment workers in Cambodia range from withheld wages to forced alcohol use, unwanted touching, verbal abuse, and physical violence [16–19]. In addition, female entertainment workers have a limited ability to exercise their rights to organize and bargain collectively for improved work conditions because of the informal nature of their work [20]. There are few or no legal channels to voice grievances or advocate for their interests because the entertainment and sex industries are mostly unregulated environments. With the passing of Cambodia’s Law on the Suppression of Human Trafficking and Sexual Exploitation in 2008 (“the trafficking law”), brothels and sex work have been criminalized, and many women moved to entertainment venues to sell sex. Police now regularly raid entertainment venues and harass or arrest female entertainment workers for selling sex [21]. Despite the Ministry of Interior of Cambodia issuing a Directive, which condoms would not be used as evidence for arrest in 2011, police continue to use the possession of condoms as evidence that someone is selling sex during raids [21, 22]. The criminalization of sex work has created conditions, where sex workers are deterred from carrying condoms, have less power to negotiate condom use [23], and are more exposed to violence from both clients and law enforcement [11, 21, 22]

The *Mobile Link* intervention is a mobile health (mHealth) project that is engaging female entertainment workers through short message service (SMS) and voice messages (VM) and linking them to the existing prevention, care and treatment services in the country. The details of the *Mobile Link* intervention and the trial design have been published elsewhere [24]. The purpose of this baseline study is to understand more about the lives of female entertainment workers in Cambodia. It provides baseline findings from the *Mobile Link* trial on the childhood conditions, pathways to sex work, HIV risk perception, contraception use, and experiences with gender-based violence of female entertainment workers in Cambodia.

## Materials and Methods

### Ethics statement

This study was approved by the National Ethics Committee for Health Research (No. 142NECHR) within the Ministry of Health in Cambodia and the Touro College Institutional Review Board (No. PH-0117). A written informed consent was obtained from each respondent. In addition, all key personnel involved in this study completed the online research ethics course on the protection of human research participants of the National Institute of Health. We acknowledged that this study required asking participants personal information about sensitive topics. We offered all participants escorted referrals to peer counselors and required services upon request.

### Settings and sampling

Data used for this study were collected as part of the baseline survey of the *Mobile Link* in March 2018 in the capital city of Phnom Penh and three other provinces Banteay Meanchey, Battambang and Siem Reap. We used a stratified random sampling method to recruit study participants. We selected entertainment venues from a list of all entertainment venues in the study sites based on a recent report on geographic information system mapping of HIV key populations in Cambodia [25]. Venues were matched with 30 similar venues and then randomized for size and type of the venues. At the venues, we approached all female entertainment workers who worked at that facility. We continued to sample from the list of all entertainment venues until we had a total of 600 female entertainment workers recruited.

Female data collectors underwent two days of training on data collection procedures. They worked in pairs and approached each selected participant to invite them to participate in the trial. If they agreed, they were asked questions regarding the inclusion/exclusion criteria. Peer data collectors asked each potential participant questions to determine if they met the nine eligibility criteria: (1) live and work at an entertainment venue in Cambodia, (2) currently sexually active defined as having engaged in oral, vaginal, or anal sex in the past three months, (3) own their own mobile phone, (4) know how to retrieve voice massages (VM) or retrieve and read short message service (SMS) on mobile phone (Khmer or Khmer with English alphabet), (5) self-identify as a female entertainment worker, (6) work in an entertainment venue and work in entertainment section at the venue, (7) were willing to receive at most one SMS/VM per day for one year, (8) provide a written informed consent, and (9) agree to a follow-up visit at six months and one year after the commencement of the intervention..

For those who met the criteria, recruiters explained the details of the study and asked them for their informed consent to participate in the study. The peer data collector verbally explained the study by reading the participant information provided as part of the informed consent process. If a female entertainment worker wished to participate, they would sign or provide their thumbprint on two copies of the consent form and were given one to keep.

### Data collection

The questionnaire contained 102 questions in the following categories: socio-demographic characteristics, entertainment work, condom use self-efficacy, HIV risk perception, HIV testing and treatment, STI testing and treatment, contraception and pregnancy, gender-based violence, substance use, and linkages to services. Peer data collectors used the Kobo Humanitarian Toolbox installed in Android operating Tablet, an open-source field data collection software on tablets, to collect the data. The interview was conducted at various places within the study sites based on agreement from individual FEW via contact by field researchers prior to the interview. The *Mobile Link* field staff took part in the study by supporting in making appointment, follow up appointment, and guiding to location. The interview took about 25 minutes using a structured Khmer questionnaire. The quality check and technical support were done during the field work by a research assistant and a research fellow. This electronic data collection system was employed to reduce data entry errors in the field. Participants were offered US$5 for time and transport compensation upon the completion of the interview.

### Data management and analyses

Survey responses that were entered into the software and exported into STATA 15 (StataCorps 2017) for statistical analyses. All variables were analyzed using descriptive statistics including means for continuous variables and proportions for categorical variables. Tables were created for each domain including demographic characteristics and childhood living conditions, pathway to entertainment work, sexual behavior and condom use, HIV risk and risk perception, contraceptive use and pregnancy experience, gender-based violence, and linkage to services.

## Results

### Demographic characteristics and childhood living conditions

As shown in Table 1, the average age of the participants was 24.3 years (SD= 3.72), and 64.5% were born in rural areas. Most participants had both parents still living (57.4%) or one parent still living (36.5%). Childhood homes had either iron/aluminum roofing (42.2%) or thatched roofs (41.8%). Fewer had ceramic or tiled roofs (14.0%). Most participants did not have piped water (87.0%) or electricity (82.0%) in their childhood home. Most had wood or bamboo plank flooring (86.2%). In terms of food security, 53.5% of participants reported that they often did not have enough food in childhood, and 14.7% said their family could not afford to send them to school. Participants reported completing primary school (46.0%), secondary school (39.7%), high school (12.5%), and university education (1.8%).

**Table 1.**
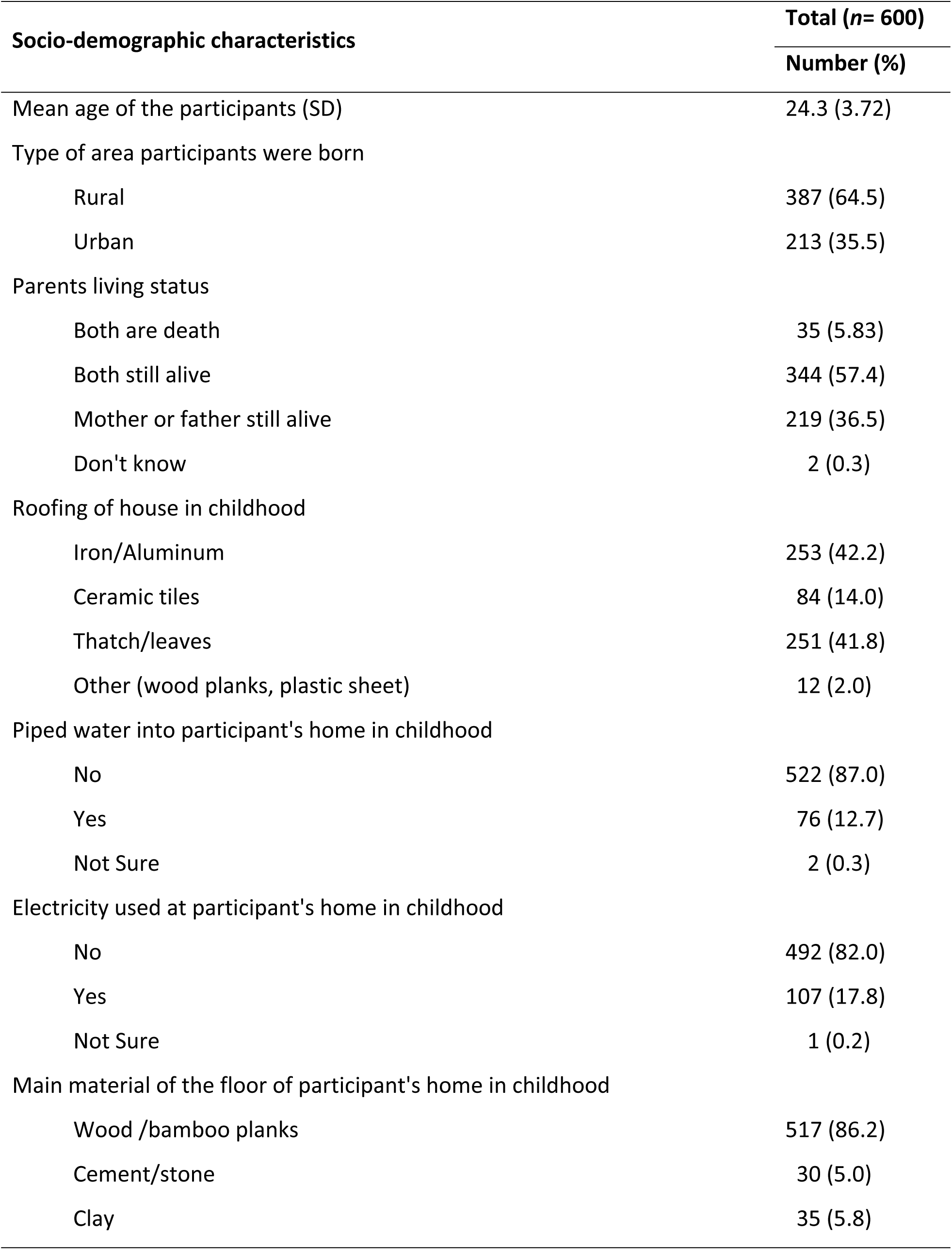
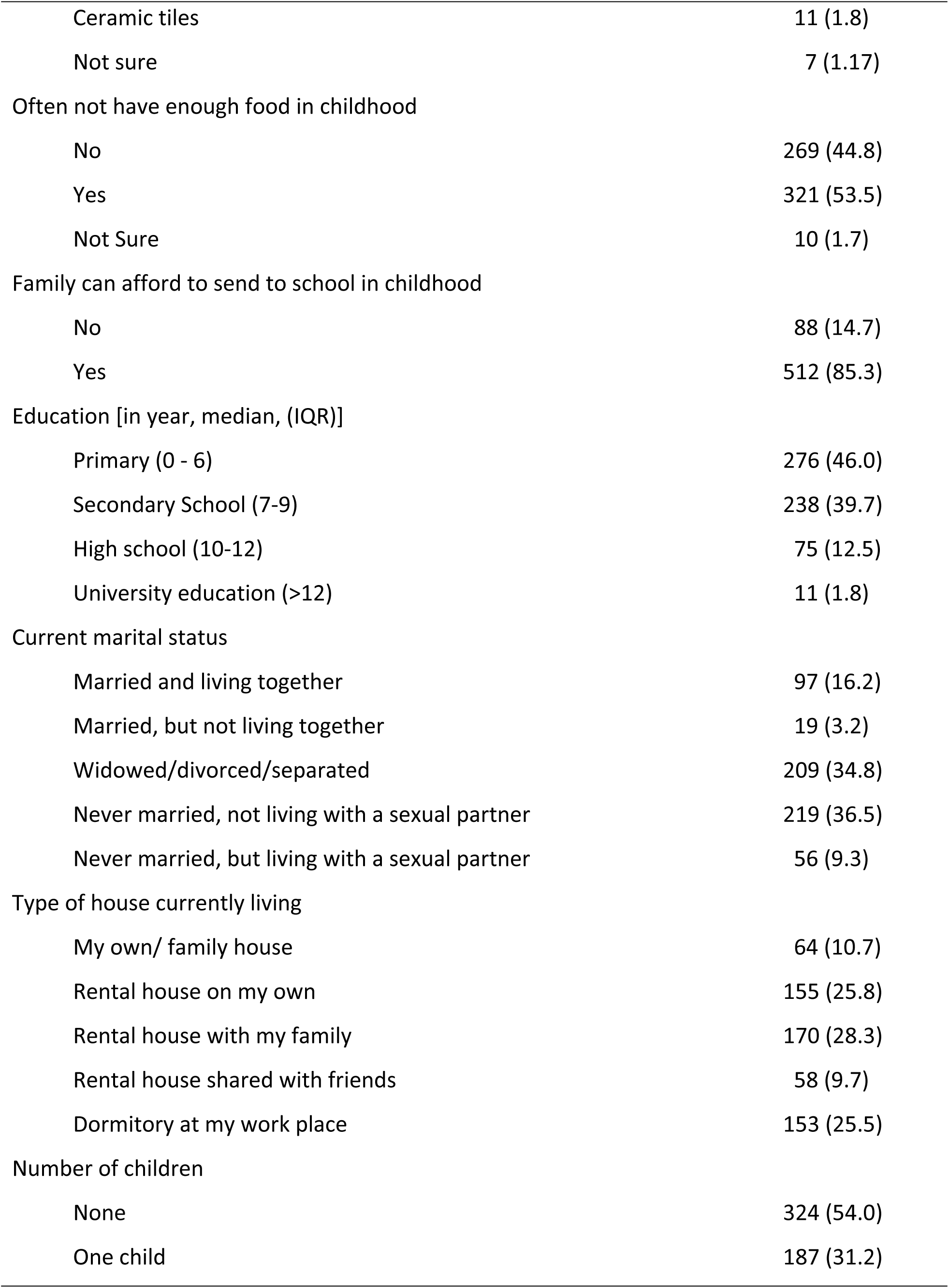
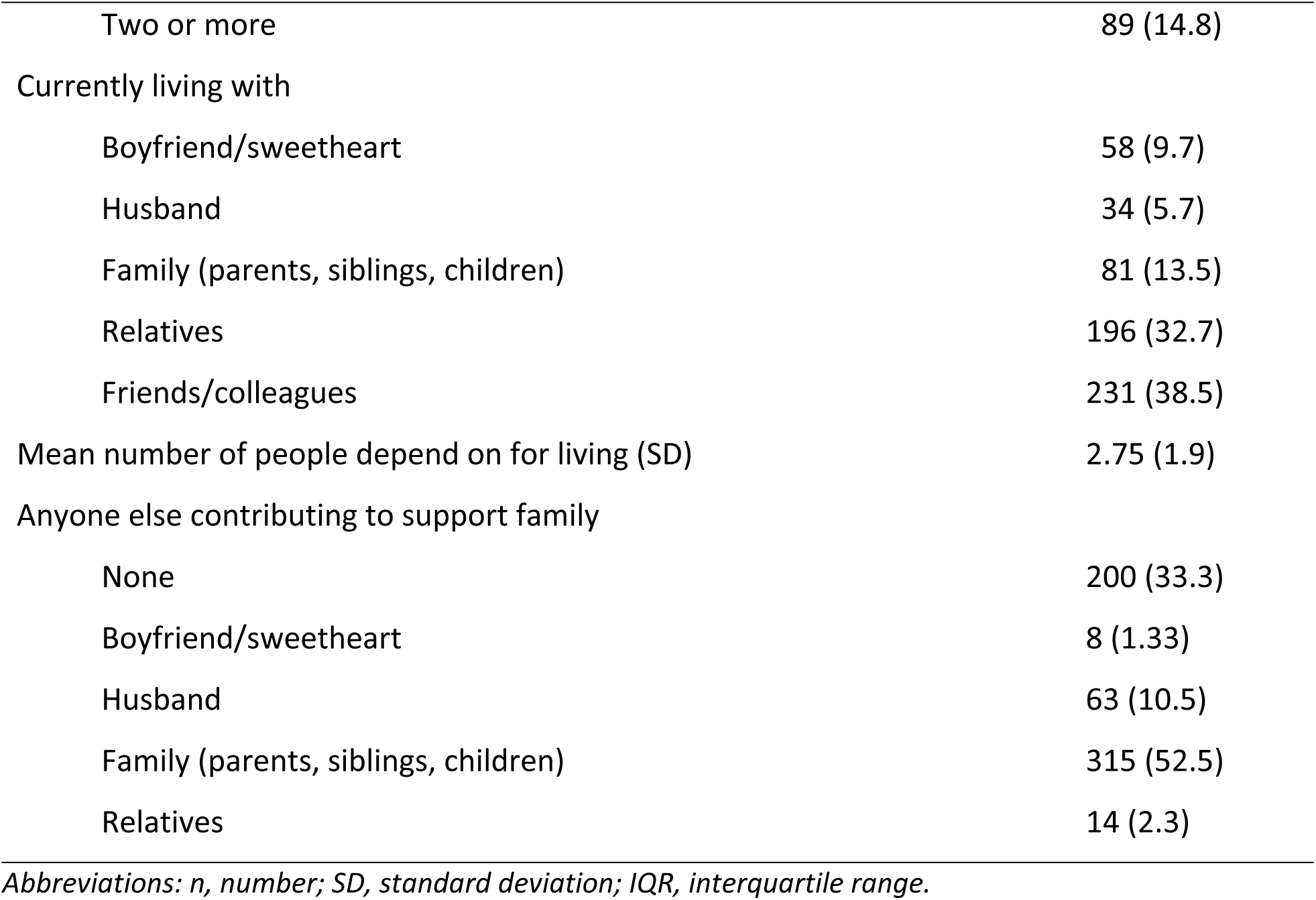
Socio-demographic characteristics of female entertainment workers

| **Socio-demographic characteristics** | **Total (n= 600)** |
| --- | --- |
|  | **Number (%)** |
| Mean age of the participants (SD) | 24.3 (3.72) |
| Type of area participants were born |  |
| Rural | 387 (64.5) |
| Urban | 213 (35.5) |
| Parents living status |  |
| Both are death | 35 (5.83) |
| Both still alive | 344 (57.4) |
| Mother or father still alive | 219 (36.5) |
| Don't know | 2 (0.3) |
| Roofing of house in childhood |  |
| Iron/Aluminum | 253 (42.2) |
| Ceramic tiles | 84 (14.0) |
| Thatch/leaves | 251 (41.8) |
| Other (wood planks, plastic sheet) | 12 (2.0) |
| Piped water into participant's home in childhood |  |
| No | 522 (87.0) |
| Yes | 76 (12.7) |
| Not Sure | 2 (0.3) |
| Electricity used at participant's home in childhood |  |
| No | 492 (82.0) |
| Yes | 107 (17.8) |
| Not Sure | 1 (0.2) |
| Main material of the floor of participant's home in childhood |  |
| Wood /bamboo planks | 517 (86.2) |
| Cement/stone | 30 (5.0) |
| Clay | 35 (5.8) |
| Ceramic tiles | 11 (1.8) |
| Not sure | 7 (1.17) |
| Often not have enough food in childhood |  |
| No | 269 (44.8) |
| Yes | 321 (53.5) |
| Not Sure | 10 (1.7) |
| Family can afford to send to school in childhood |  |
| No | 88 (14.7) |
| Yes | 512 (85.3) |
| Education [in year, median, (IQR)] |  |
| Primary (0 - 6) | 276 (46.0) |
| Secondary School (7-9) | 238 (39.7) |
| High school (10-12) | 75 (12.5) |
| University education (>12) | 11 (1.8) |
| Current marital status |  |
| Married and living together | 97 (16.2) |
| Married, but not living together | 19 (3.2) |
| Widowed/divorced/separated | 209 (34.8) |
| Never married, not living with a sexual partner | 219 (36.5) |
| Never married, but living with a sexual partner | 56 (9.3) |
| Type of house currently living |  |
| My own/ family house | 64 (10.7) |
| Rental house on my own | 155 (25.8) |
| Rental house with my family | 170 (28.3) |
| Rental house shared with friends | 58 (9.7) |
| Dormitory at my work place | 153 (25.5) |
| Number of children |  |
| None | 324 (54.0) |
| One child | 187 (31.2) |
| Two or more | 89 (14.8) |
| Currently living with |  |
| Boyfriend/sweetheart | 58 (9.7) |
| Husband | 34 (5.7) |
| Family (parents, siblings, children) | 81 (13.5) |
| Relatives | 196 (32.7) |
| Friends/colleagues | 231 (38.5) |
| Mean number of people depend on for living (SD) | 2.75 (1.9) |
| Anyone else contributing to support family |  |
| None | 200 (33.3) |
| Boyfriend/sweetheart | 8 (1.33) |
| Husband | 63 (10.5) |
| Family (parents, siblings, children) | 315 (52.5) |
| Relatives | 14 (2.3) |

*Abbreviations: n, number; SD, standard deviation; IQR, interquartile range.*

Participants reported having never been married and not living with a partner (36.5%), being widowed/divorced or separated (34.8%), being married and living together (16.2%), being never married but living with partner (9.3%), and being married but not living together (3.2%). They reported living in a rental house with their family (28.3%), on their own in a rental house (25.8%), at a dormitory at their workplace (25.5%), in a family house that they own (10.7%), or in a rental house shared with friends (9.7%). Participants were currently living with friends or colleagues (38.5%), relatives (32.7%), family (13.5%), boyfriends (9.7%), or their husband (5.7%). More than half of the participants (54.0%) had no children, while 31.2% had one child, and 14.8% had two or more children. They reported having an average of 2.8 people depending on them for living, and 52.5% reported having additional support from family.

### Pathway to entertainment work

As shown in Table 2, participants have lived in the city for one to three years (50.5%), four to 10 years (30%), 19 to 30 years (12.5%), and 11 to 18 years (7.3%). Top reasons for moving to their current city were economic opportunity (43.7%), getting away from a bad situation (9.0%), following family members (6.3%), and interest in new places (4.2%). About one-third (36.7%) reported having worked in the garment industry and that their main reasons for leaving the factory were seeking better work conditions elsewhere (21.8%), seeking better pay elsewhere (12.7%), health problems (12.3%), getting fired or laid off (11.4%), getting offered another job (10.9%), and being interested in getting a different job (6.4%).

**Table 2.** Entertainment work experiences of female entertainment works

| Entertainment work experiences | Total (n= 600) |
| --- | --- |
|  | Number (%) |
| Ever worked in the garment industry | 220 (36.7) |
| Main reason for leaving the garment industry |  |
| Better pay elsewhere | 20 (12.7) |
| Better working conditions elsewhere | 48 (21.8) |
| Was fired or laid off | 25 (11.4) |
| Interested in a different job | 14 (6.4) |
| Offered another job | 24 (10.9) |
| Had health problem | 27 (12.3) |
| Other (far from home, family reasons) | 54 (24.5) |
| Mean duration (in month) working in the entertainment industry (SD) | 23.6 (25.4) |
| Type of venue best describes first job in entertainment |  |
| Karaoke bar | 219 (53.2) |
| Massage parlor | 10 (1.7) |
| Beer garden | 54 (9.0) |
| Restaurant/cafe | 204 (34.0) |
| Dance club | 11 (1.8) |
| Freelance (street, public parks) | 2 (0.3) |
| Type of venue best describes current job in entertainment |  |
| Karaoke bar | 357 (59.5) |
| Massage parlor | 14 (2.3) |
| Beer garden | 50 (8.3) |
| Restaurant/cafe | 175 (29.2) |
| Freelance (street, public parks) | 4 (0.7) |
| Income per month from current entertainment job [USD, median (IQR)] | 200 (150) |
| < 100 | 24 (4.0) |
| 100 - 199 | 167 (27.8) |
| 200 - 299 | 221 (36.8) |
| > 300 | 188 (31.3) |
| Part of any organization that support entertainment/ sex workers | 38 (6.3) |

*Abbreviations: SD, standard deviation; IQR, interquartile range.*

On average, they have worked in the entertainment industry for 23.6 months (SD=25.4). The first venue they worked for was karaoke bars (53.2%), restaurants or cafes (34%), beer gardens (9%), massage parlors (1.9%), and dance clubs (1.8%). Only 0.3% were freelance or street-based entertainment workers. They were currently working in karaoke bars (59.5%), restaurants or cafes (29.2%), beer gardens (8.3%), massage parlors (2.3%), and as freelancers (0.7%). Reported median monthly income in the past six months was US$200 USD (IQR=150), with 36.8% reporting the income in the US$200-299 range, 31.3% over US$300, and 27.8% in the US$100-199 range. Only 6.3% reported that they were part of an organization that support FEWs.

### Sexual behaviors, condom use, and HIV testing

Table 3 shows that 65.3% of participants reported having had sexual intercourse, and 76.3% had sex not in exchange for money or gifts in the past three months. The mean number of sexual partners not in exchange for money or gifts was 1.1 (SD= 0.3), with 95.3% reporting having one partner not in exchange for money or gifts in the past three months. Regarding condom use, 23.4% reported using a condom at the last sexual intercourse, and 68.2% reported never using a condom with this type of sexual partner in the past three months. The main reasons for not using a condom with a partner not in exchange for money or gifts were that they trusted their partners (43.8%), they did not like using condoms (13.4%), their partners requested not to use condoms (12.4%), or they did not think about it (11.3%).

**Table 3.**
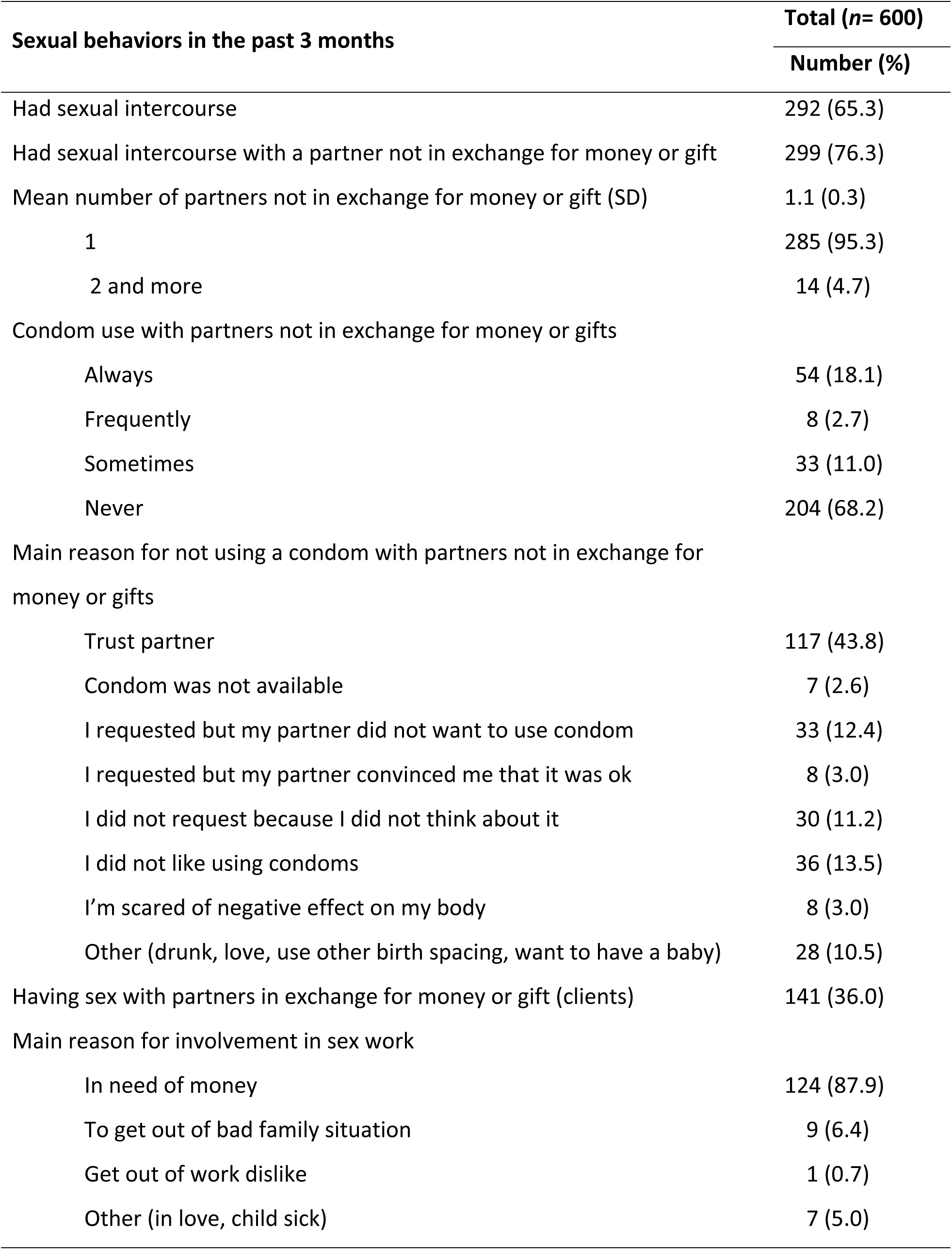
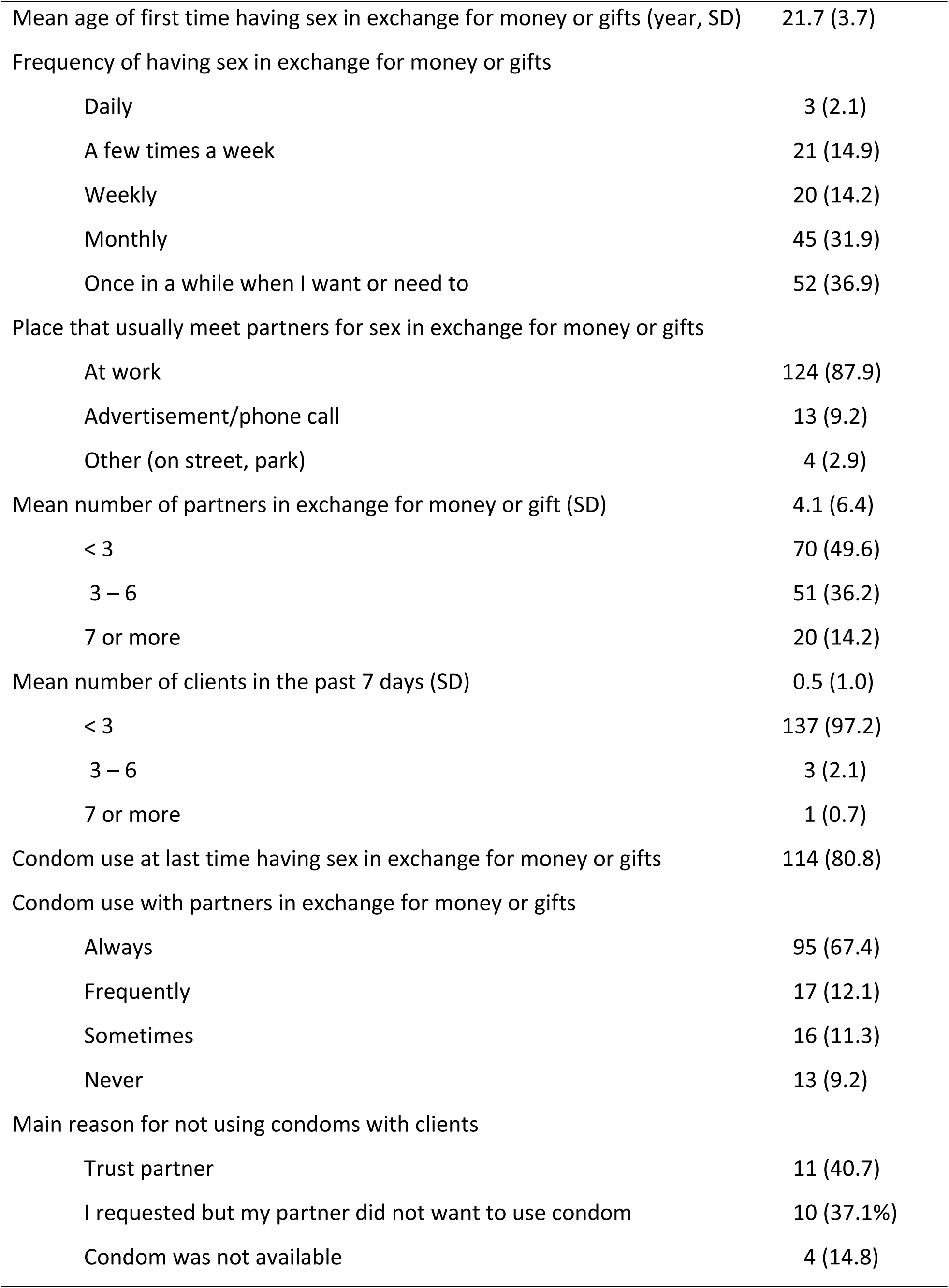
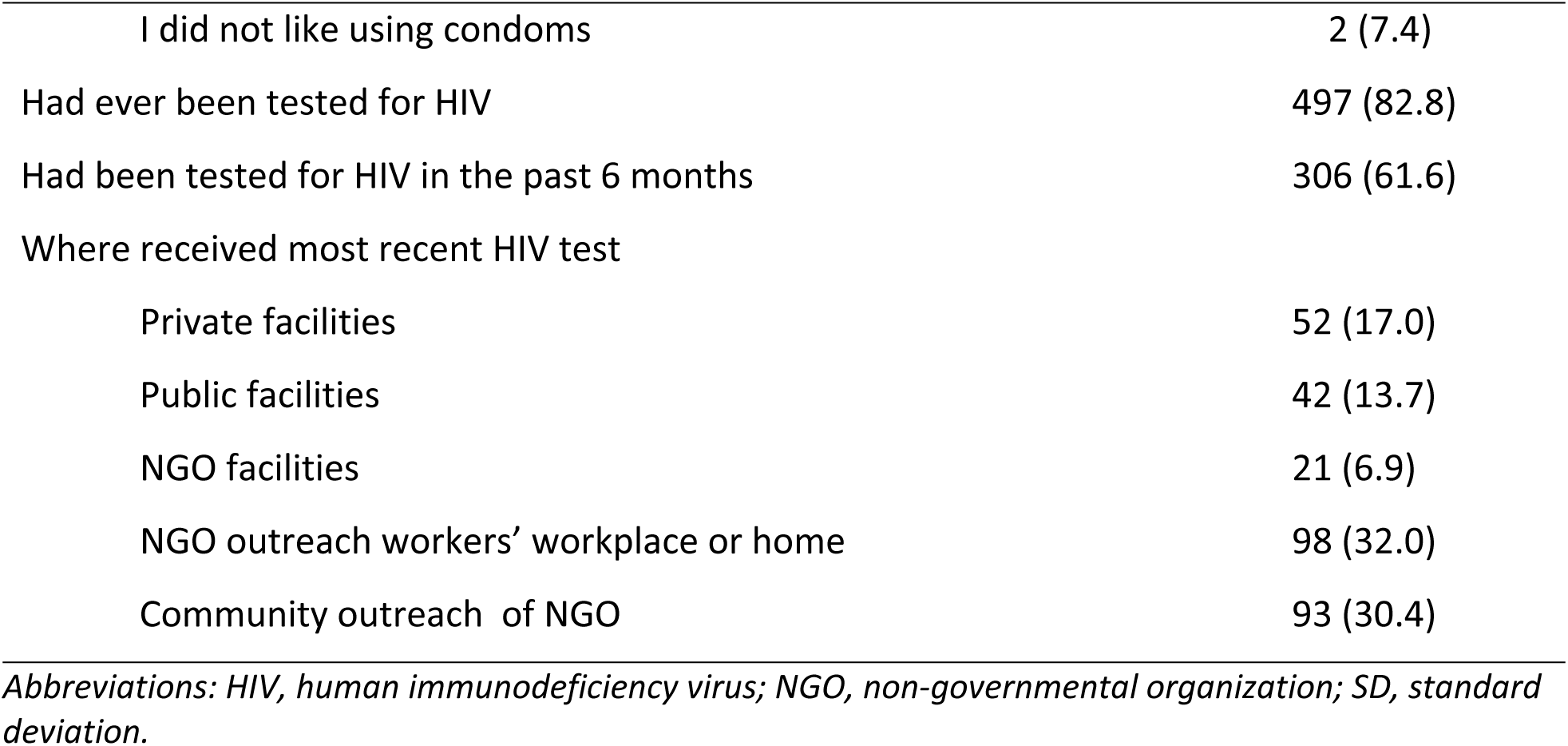
Sexual behaviors, condom use, and HIV testing of female entertainment workers

| Sexual behaviors in the past 3 months | Total (n= 600) |
| --- | --- |
|  | Number (%) |
| Had sexual intercourse | 292 (65.3) |
| Had sexual intercourse with a partner not in exchange for money or gift | 299 (76.3) |
| Mean number of partners not in exchange for money or gift (SD) | 1.1 (0.3) |
| 1 | 285 (95.3) |
| 2 and more | 14 (4.7) |
| Condom use with partners not in exchange for money or gifts |  |
| Always | 54 (18.1) |
| Frequently | 8 (2.7) |
| Sometimes | 33 (11.0) |
| Never | 204 (68.2) |
| Main reason for not using a condom with partners not in exchange for money or gifts |  |
| Trust partner | 117 (43.8) |
| Condom was not available | 7 (2.6) |
| I requested but my partner did not want to use condom | 33 (12.4) |
| I requested but my partner convinced me that it was ok | 8 (3.0) |
| I did not request because I did not think about it | 30 (11.2) |
| I did not like using condoms | 36 (13.5) |
| I'm scared of negative effect on my body | 8 (3.0) |
| Other (drunk, love, use other birth spacing, want to have a baby) | 28 (10.5) |
| Having sex with partners in exchange for money or gift (clients) | 141 (36.0) |
| Main reason for involvement in sex work |  |
| In need of money | 124 (87.9) |
| To get out of bad family situation | 9 (6.4) |
| Get out of work dislike | 1 (0.7) |
| Other (in love, child sick) | 7 (5.0) |
| Mean age of first time having sex in exchange for money or gifts (year, SD) | 21.7 (3.7) |
| Frequency of having sex in exchange for money or gifts |  |
| Daily | 3 (2.1) |
| A few times a week | 21 (14.9) |
| Weekly | 20 (14.2) |
| Monthly | 45 (31.9) |
| Once in a while when I want or need to | 52 (36.9) |
| Place that usually meet partners for sex in exchange for money or gifts |  |
| At work | 124 (87.9) |
| Advertisement/phone call | 13 (9.2) |
| Other (on street, park) | 4 (2.9) |
| Mean number of partners in exchange for money or gift (SD) | 4.1 (6.4) |
| < 3 | 70 (49.6) |
| 3 – 6 | 51 (36.2) |
| 7 or more | 20 (14.2) |
| Mean number of clients in the past 7 days (SD) | 0.5 (1.0) |
| < 3 | 137 (97.2) |
| 3 – 6 | 3 (2.1) |
| 7 or more | 1 (0.7) |
| Condom use at last time having sex in exchange for money or gifts | 114 (80.8) |
| Condom use with partners in exchange for money or gifts |  |
| Always | 95 (67.4) |
| Frequently | 17 (12.1) |
| Sometimes | 16 (11.3) |
| Never | 13 (9.2) |
| Main reason for not using condoms with clients |  |
| Trust partner | 11 (40.7) |
| I requested but my partner did not want to use condom | 10 (37.1%) |
| Condom was not available | 4 (14.8) |
| I did not like using condoms | 2 (7.4) |
| Had ever been tested for HIV | 497 (82.8) |
| Had been tested for HIV in the past 6 months | 306 (61.6) |
| Where received most recent HIV test |  |
| Private facilities | 52 (17.0) |
| Public facilities | 42 (13.7) |
| NGO facilities | 21 (6.9) |
| NGO outreach workers' workplace or home | 98 (32.0) |
| Community outreach of NGO | 93 (30.4) |

*Abbreviations: HIV, human immunodeficiency virus; NGO, non-governmental organization; SD, standard deviation.*

Of the total participants, 36.0% reported having had sexual intercourse with a partner in exchange for money or gifts (client) in the past three months. The main reasons for involvement in sex work was that they needed money (87.9%), to get out of bad family situation (6.4%), or they had a child getting sick (5.0%). The mean age of first involvement in sex work was 21.7 years old (SD= 3.7). Frequency of involvement in sex work was once in a while when they wanted or needed to (36.9%), monthly (31.9%), a few times a week (14.9%), weekly (14.2%), and daily (2.1%). Participants described meeting clients at work (87.9%), through advertisements (9.2%), and on the street or in the parks (2.9%). The mean number of clients in the past three months was 4.1 (SD= 6.4), with 49.6% reporting less than three clients, 36.2% reporting three to six clients, and 14.2% reporting seven or more clients. The majority (80.8%) reported using a condom at last sex with client. More than two-thirds (67.4%) reported always using condoms with clients in the past three months. The main reasons for not using condoms with clients included trust (40.7%), clients did not want to use (37.1%), condom was not available (14.8%), and they did not like condoms (7.4%). Overall, 61.6% of participants reported having been tested for HIV in the past six months. The places where they were tested for HIV was at an NGO outreach workers’ workplace or home (32.0%), a community outreach program of NGO (30.4%), at a private health facility (17.0%), at a public health facility (13.7%), and at an NGO facility (6.9%). All respondents reported that the result of their most recent HIV test was negative.

### Contraceptive use and pregnancy experience

Table 4 shows that 31.4% reported using modern contraception including condoms (37.4%), pills (41.2%), injectable (11.2%), implant (3.2%), intrauterine devices (2.7%), and other methods such as sterilization and emergency contraception (1.7%). Reasons for not using a modern contraception were that they did not think they needed contraception (55.6%), did not want to prevent pregnancy (13.3%), did not like the side effects of contraception (10.9%), did not like using modern methods (5.1%), and other reasons such as health issues, no sex partner, or use of calendar method (12.1%), or they did not know where to get contraception (1.8%).

**Table 4.** Contraceptive use and pregnancy experience of female entertainment works

| **Contraceptive use and pregnancy** | **Total (n= 600)** |
| --- | --- |
|  | **Number (%)** |
| Currently use modern contraceptive to prevent pregnancy | 188 (31.4) |
| Type of contraceptive using |  |
| Male condom | 70 (37.4) |
| Pill | 77 (41.2) |
| Injectable | 21 (11.2) |
| Intra-uterus devices (IUD) | 5 (2.7) |
| Implant (use under the skin) | 6 (3.2) |
| Female sterilization | 3 (1.6) |
| Other (male sterilization, use 72-hour pill) | 5 (2.7) |
| Reasons of not currently using any modern contraceptive method |  |
| Do not like side effects | 36 (10.9) |
| Do not think I need contraception | 184 (55.6) |
| Do not want to prevent pregnancy/want to get pregnant | 44 (13.3) |
| Do not like using modern methods | 17 (5.1) |
| Do not know where to get contraception | 6 (1.8) |
| Do not think I can afford contraception | 4 (1.2) |
| Other (health issue, no partner, calendar method, no partner) | 40 (12.1) |
| Thought medical abortion (before 12 weeks gestation) is legal in Cambodia |  |
| No | 378 (63.0) |
| Yes | 100 (16.7) |
| Don't know | 122 (20.4) |
| Experienced unwanted pregnancy in lifetime | 265 (44.2) |
| Experienced unwanted pregnancy in the past 12 months | 97 (36.6) |
| Had an abortion in lifetime | 210 (79.2) |
| Mean number of abortion in life time (SD) | 1.7 (1.3) |
| 1 time | 135 (64.3) |
| 2 to 3 | 62 (29.5) |
| 4 or more | 13 (6.2) |
| Mean number of abortion in the past 12 months (SD) | 0.5 (0.6) |
| None | 127 (60.5) |
| 1 time | 72 (34.3) |
| 2 or more | 11 (5.2) |
| Facility getting service for the most recent abortion |  |
| Medication from a pharmacy | 127 (60.5) |
| Private facility | 66 (31.4) |
| Public facility | 8 (3.8) |
| NGO facility | 9 (4.3) |
| Experience complications from most recent abortion | 58 (27.6) |
| Seek treatment for the complications | 41 (70.7) |
| Facilities seeking treatment for the complications |  |
| Medication from a pharmacy | 7 (17.1) |
| Private facility | 18 (43.9) |
| Public facility | 7 (17.1) |
| NGO facility | 4 (9.8) |
| Traditional healer | 3 (7.3) |
| Other (call mobile practitioner, self-care) | 2 (4.8) |

*Abbreviations: NGO, non-governmental organization; SD, standard deviation.*

Of all participants, 63.0% thought medical abortion was not legal in Cambodia, and 36.6% reported having an unwanted pregnancy in the past 12 months. A vast majority of the participants (79.2%) reported ever having an induced abortion with a mean number of lifetime abortions of 1.7 (SD= 1.3), and a mean number of abortions in last 12 months of 0.5 (SD= 0.6). Participants reported getting abortion services for the most recent abortion at pharmacies (60.5%), private clinics (31.4%), NGO clinics (4.3%), and public clinics (3.8%). Of those who had an abortion, 27.6% reported experiencing a complication such as excessive bleeding or infection, and 70.7% reported seeking treatment for these complications from private clinics (43.9%), pharmacies (17.1%), public clinics (17.1%), NGO clinics (9.8%), traditional healers (7.3%), or other facilities (4.8%).

### Gender-based violence

As shown in Table 5, 21.8% of the participants reported experiencing unwanted touching or groping at work in the past three months. In response to the abuse, 33.2% felt that there is nothing to do, 29.8% would tell other family members or friends, 16.2% would go to the police or the court, 9.8% would go to an NGO, and 2.3% would go to local authorities. In terms of justification of abuse, 82.5% of participants said that physical abuse was not justified when a wife does not obey her husband, 60% said that verbal abuse was not justified when a wife does not obey her husband, and 87.2% said that physical abuse was not justified when a girlfriend does not obey her boyfriend. Over half (64%) of the participants reported lifetime experiences with gender-based violence including forced use of alcohol (23.0%), physical abuse (19.2%), verbal threats (13.0%), controlling ability to leave the house (2.8%), and forced sex (2.0%). Just less than half of the participants (45.4%) reported experiences with gender-based violence in the past six months. Of those who had experienced gender-based violence, participants reported experiencing forced use of alcohol (20.9%), verbal threats (14.4%), physical abuse (6.7%), and forced sex (2.1%). The main perpetrators of the reported violence included clients (54.3%), husband/partners (22.6%), other family members, friends, strangers, taxi drivers (18.6%), and sweethearts (5.5%).

**Table 5.**
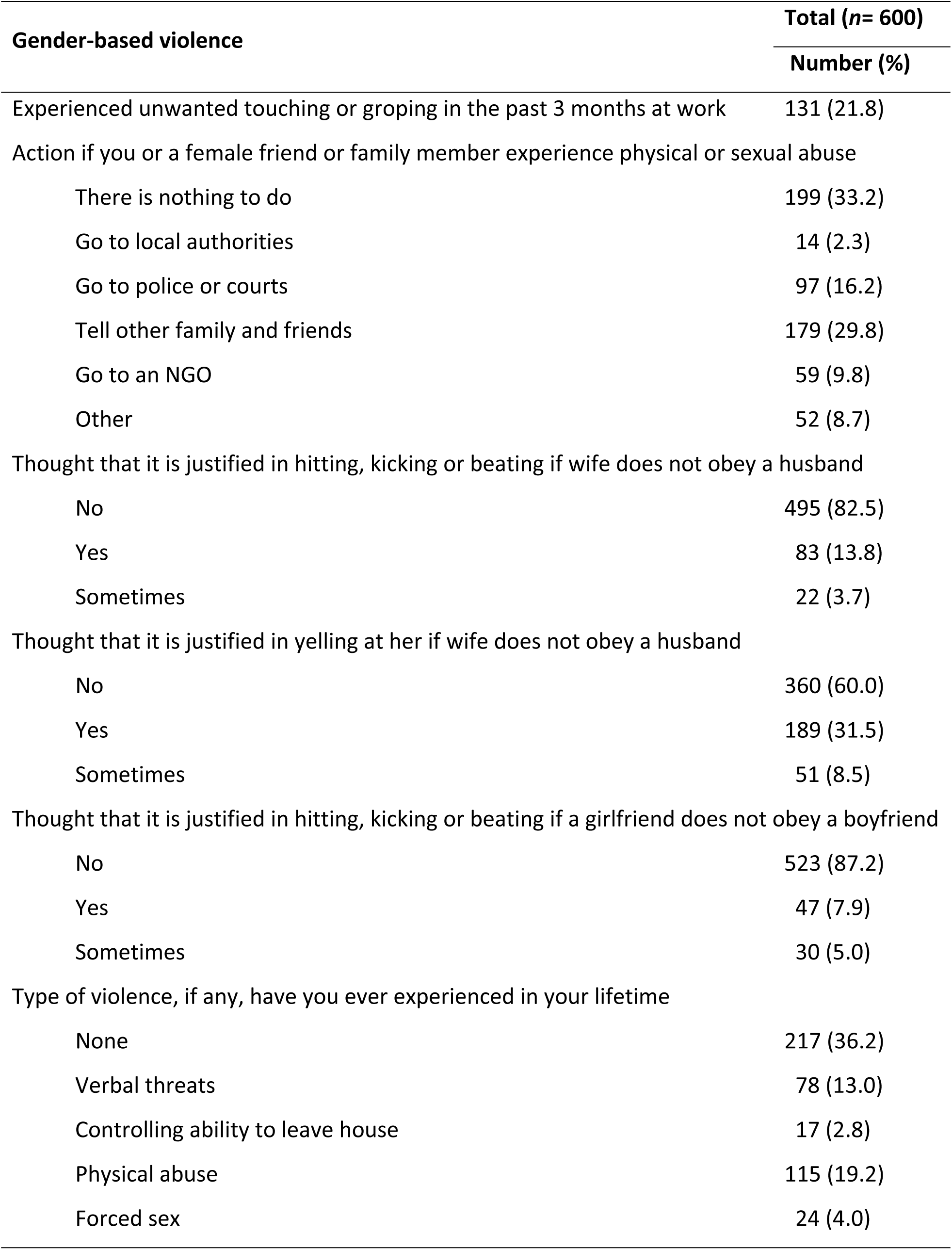
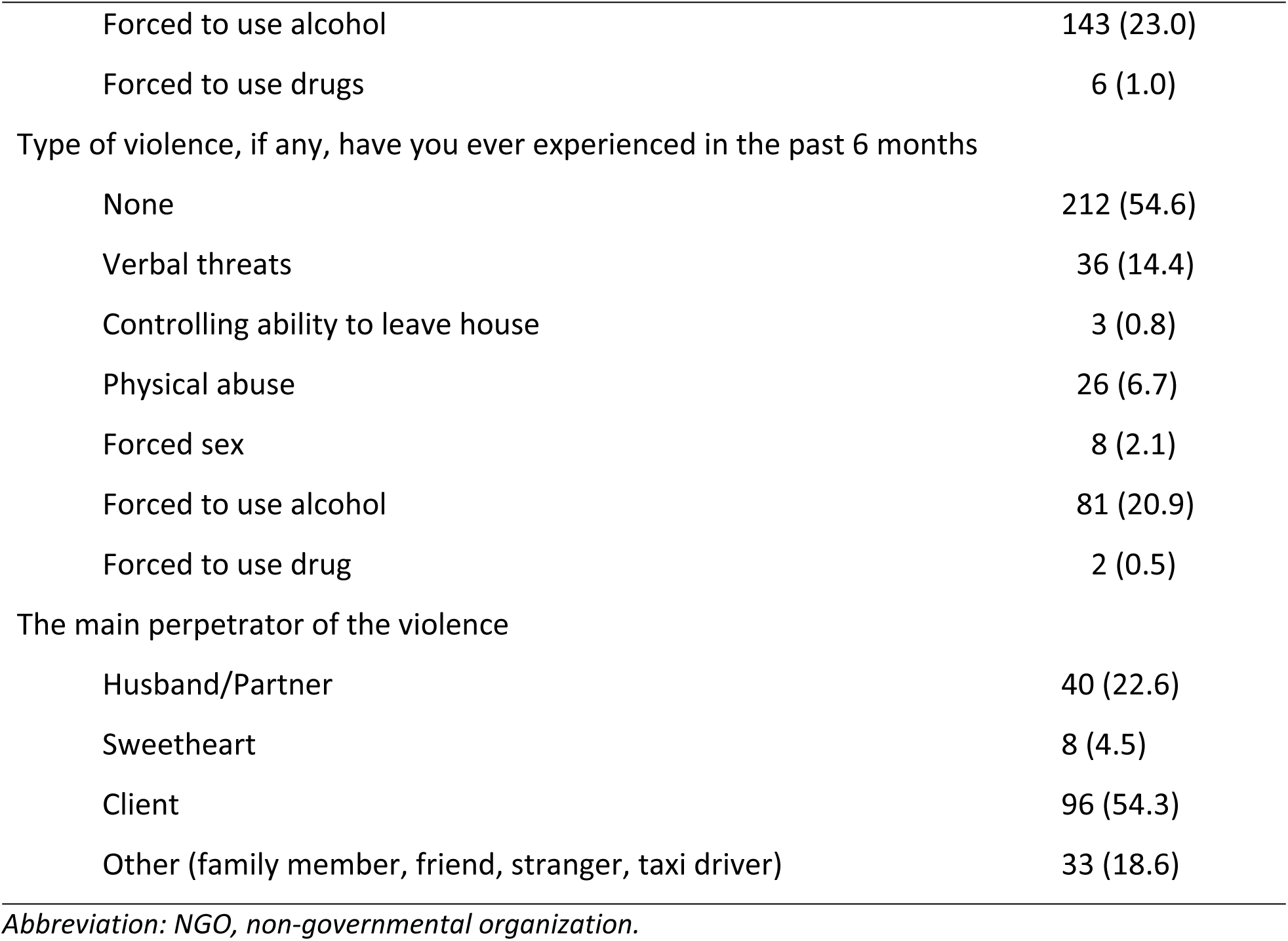
Gender-based violence among female entertainment workers

| **Gender-based violence** | **Total (n= 600)** |
| --- | --- |
|  | **Number (%)** |
| Experienced unwanted touching or groping in the past 3 months at work | 131 (21.8) |
| Action if you or a female friend or family member experience physical or sexual abuse |  |
| There is nothing to do | 199 (33.2) |
| Go to local authorities | 14 (2.3) |
| Go to police or courts | 97 (16.2) |
| Tell other family and friends | 179 (29.8) |
| Go to an NGO | 59 (9.8) |
| Other | 52 (8.7) |
| Thought that it is justified in hitting, kicking or beating if wife does not obey a husband |  |
| No | 495 (82.5) |
| Yes | 83 (13.8) |
| Sometimes | 22 (3.7) |
| Thought that it is justified in yelling at her if wife does not obey a husband |  |
| No | 360 (60.0) |
| Yes | 189 (31.5) |
| Sometimes | 51 (8.5) |
| Thought that it is justified in hitting, kicking or beating if a girlfriend does not obey a boyfriend |  |
| No | 523 (87.2) |
| Yes | 47 (7.9) |
| Sometimes | 30 (5.0) |
| Type of violence, if any, have you ever experienced in your lifetime |  |
| None | 217 (36.2) |
| Verbal threats | 78 (13.0) |
| Controlling ability to leave house | 17 (2.8) |
| Physical abuse | 115 (19.2) |
| Forced sex | 24 (4.0) |
| Forced to use alcohol | 143 (23.0) |
| Forced to use drugs | 6 (1.0) |
| Type of violence, if any, have you ever experienced in the past 6 months |  |
| None | 212 (54.6) |
| Verbal threats | 36 (14.4) |
| Controlling ability to leave house | 3 (0.8) |
| Physical abuse | 26 (6.7) |
| Forced sex | 8 (2.1) |
| Forced to use alcohol | 81 (20.9) |
| Forced to use drug | 2 (0.5) |
| The main perpetrator of the violence |  |
| Husband/Partner | 40 (22.6) |
| Sweetheart | 8 (4.5) |
| Client | 96 (54.3) |
| Other (family member, friend, stranger, taxi driver) | 33 (18.6) |

*Abbreviation: NGO, non-governmental organization.*

### Alcohol use

Table 6 shows that 71.0% of the participants reported having at least one standard alcoholic drink four or more times per week (14.5% having one drink 2-3 times per week, 4.3% had 2-4 times per month, and 6.7% having once a month or less). Regarding the amount of alcohol, 31.7% reported having 10 or more standard alcoholic drinks, 5.9% having 7-9 standard alcoholic drinks, 27.3% having 5-6 standard alcoholic drinks, 20.4% having 3-4 standard alcoholic drinks, 14.7% having 1-2 standard alcoholic drinks of alcohol on a typical day they drank in the past three months. About one in five participants (20.1%) reported having been forced to drink alcohol at least once a week in the past three months.

**Table 6.**
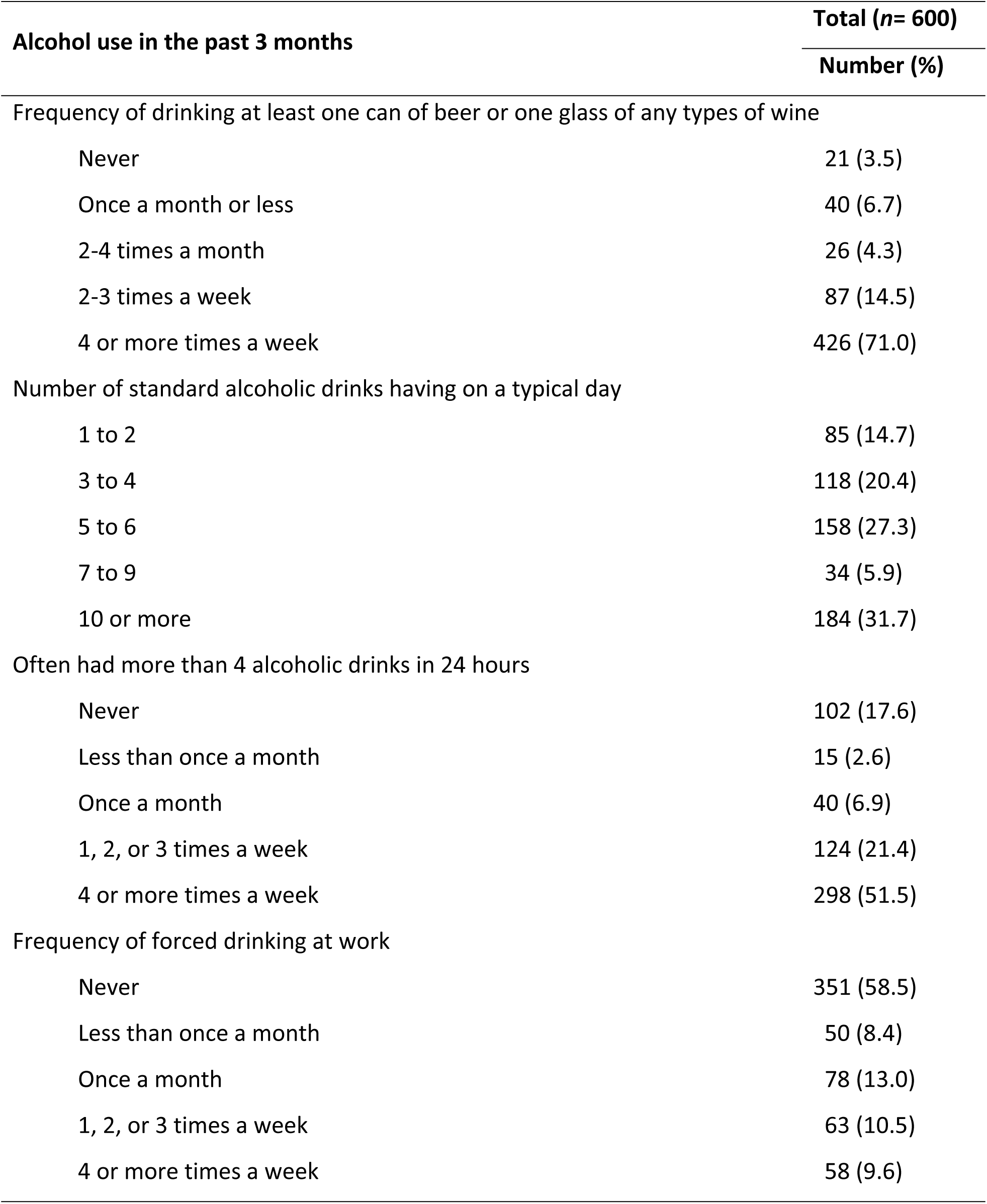
Alcohol use among female entertainment workers

### Linkage to services

Table 7 shows that 20% of the participants reported having contacted an outreach worker to ask a health question in the last six months. Health issues they contacted about were vaginal health (82.5%), family planning (4.3%), HIV (4.2%), STIs (4.2%), and other concerns (5.8%). The number of times they had reached out to an outreach worker was one time (24.2%), two to four times (20.0%), and five times or more (2.5%) in the past six months. Two-thirds of the participants (60%) have received an escorted referral for a health service from an outreach worker for vaginal health (80.5%), family planning (9.7), HIV testing (5.5%), and STI testing (4.2%).

**Table 7.** Linkage to health services among female entertainment workers

| Linkage to health services in the past 6 months | Total (n= 600) |
| --- | --- |
|  | Number (%) |
| Contacted outreach workers to ask health questions | 120 (20.0) |
| Main health issues for which you have contacted outreach workers |  |
| HIV | 5 (4.2) |
| STIs | 5 (4.2) |
| Family planning | 4 (3.3) |
| Vaginal health (discharge, irritation, inflammation) | 99 (82.5) |
| Other | 9 (5.8) |
| Number of times you have contacted outreach workers |  |
| Never | 64 (53.3) |
| 1 time | 29 (24.2) |
| 2-4 times | 24 (20.0) |
| 5+ times | 3 (2.5) |
| Received an escorted referral for health services from outreach workers | 72 (60.0) |
| Health issues did you receive an escorted referral |  |
| HIV | 4 (5.5) |
| STIs | 3 (4.2) |
| Family planning | 7 (9.7) |
| Vaginal health (discharge, irritation, inflammation) | 58 (80.6) |

*Abbreviations: HIV, human immunodeficiency virus; STI, sexually transmitted infections.*

## Discussion

Several important findings are noteworthy including low condom use practices with non-paying partners, reports of excessive and forced alcohol use at work, modern contraception use, and experiences of gender-based violence.

In this study, condom use at last sex with non-paying partners was reported by 23% of respondents. This is both higher than and lower than other recent reports from FEWs. Studies among FEWs in Cambodia found that 12% and 15% of FEWs reported consistent condoms use with non-paying partners [25, 26]. A recent study conducted by the authors of this paper found that 38% reported using a condom at last intercourse with non-commercial partners [27]. There are many reasons that may explain inaccurate reporting on condom use, but all three rates suggest that the majority of FEWs have condomless sex with their non-paying partners. In our study, women stated that the main reason for not using a condom with a non-paying partner was that they trusted their partner, they do not like using condoms, their partner did not want to, or they did not think about it. In a study of female sex workers in India, the odds of consistent condom use with husbands or other non-paying partners was higher when their partner knew they engaged in sex work and if they were unmarried. Also, the longer the relationship the less likely to use condoms consistently [28].

Our study found that 33% of respondents reported being forced to drink alcohol at work more than once a month. While there is not a lot of data on reported forced drinking among FEWs, some evidence from other studies can support our finding. A study on FEWs in China found that 57% of respondents had a high score on a risky drinking measure [19]. A study of sex workers in Cambodia found that 85% self-reported unhealthy alcohol consumption [18]. A qualitative study from Cambodia [17] found that heavy drinking was high among FEWs and was considered a norm in entertainment work. In addition, women connected their alcohol use or their clients’ alcohol use to condomless sex [17]. In 2014, the Cambodian Ministry of Labor and Vocational Training extended occupational safety laws to include all entertainment workers and add specific regulations to protect against risks faced by all entertainment workers specifically, including forcing “entertainment workers to work overtime, drink alcohol, use drugs or undergo abortions” [16]. The findings from this study bring up questions about how compliance is monitored.

Of respondents in our study, 59% reported currently not using modern contraception mostly because either they wanted to become pregnant (45%) or they did not like the side effects of modern methods (30%). This contraceptive use rate is higher than the rates reported in other studies in Cambodia. Another study of 204 FEWs in Phnom Penh, Cambodia conducted between 2009 and 2010 found that 11% reported hormonal contraceptive use in the past three months [26]. Over the eight years between these two studies, many sexual and reproductive health services have worked to support the growing number of FEWs, and national contraceptive prevalence rate has been increasing [29]. These efforts may account for the significant difference between these two estimates.

In this study, 23% of FEWs reported verbal threats, physical abuse, or forced sex in past six months. In another study of FEWs in Cambodia, 48% reported physical or sexual violence in the past year [25]. They also found that experience of violence was associated with drug use and decreased the odds of consistent condom use with non-paying partners. In a related study about police violence against key populations in Cambodia, 27% of FEWS reported being verbally threatened by police, 13% reported being forced to pay money to avoid arrest, and 5% reported being forced to exchange sex to avoid arrest [21]. Our study did not ask about police violence and most respondents reported main perpetrators of violence were clients (53%) and husbands/partners (25%).

### Limitations of the study

Limitations of this descriptive study include the fact that all measures were self-reported, which may introduce several types of bias including social desirability as participants may have answered in a way that would portray them in a more positive light. Participants may have over-reported socially desirable traits (recent HIV testing) and under-reported socially undesired traits (unprotected sex with clients). In addition, recall bias could also be introduced as participants may not be able to recall their behaviors with complete accuracy, especially details such as frequency of behaviors. We asked extremely sensitive questions, and it is likely that participants could be too embarrassed to respond honestly. In addition, this study may not be representative of all FEWs in Cambodia due to sampling bias. Our sampling procedure was systematic and thorough but over-sampled from women who worked at established venues that were counted during a recent mapping project and may not have included FEWs at smaller, less established, or more covert establishments.

## Conclusions

The findings for this cross-sectional study give us a snapshot of the lives of FEWs in Cambodia today. Young women continue to migrate for work within Cambodia. With the number of FEWs in the country almost doubling in the past five years, support for their safety and health is an increasing priority. FEWs are working within a context of changing laws, regulations, and enforcement that affect their work environments. The *Mobile Link* trial is focused on individual level health information and health services access. The information that is being gathered as part of this trial will help to support the development of future individual and structural level interventions for the safety and support of FEWs as well as an evidence base that can inform policy level changes to support the realization of full human rights for entertainment works in Cambodia including the right to health, safety, and employment.

## Acknowledgements

This study was financially supported by the 5% Initiative. We thank field data collectors, staff members of our associated partners, community support volunteers, outreach workers, and participants for their contribution to data collection. We also thank Ms. Ngovlily Sok, Research Volunteer at KHANA Center for Population Health Research, for her excellent contribution to field coordination, administration, and logistic supports for this study.

